# Acetic acid is a superior ion pairing modifier for sub-nanogram and single cell proteomics

**DOI:** 10.1101/2023.08.01.551522

**Authors:** Colten D. Eberhard, Benjamin C. Orsburn

## Abstract

A recent study demonstrated a substantial increase in peptide signal and corresponding proteome coverage when employing 0.5% acetic acid (AA) as the ion pairing modifier in place of the 0.1% formic acid traditionally used in shotgun proteomics. In this study, we investigated the effect of modifier in the context of sub-nanogram and single cell proteomics (SCP). We first evaluated a tryptic digest standard down to 20 picograms total load on column on a TIMSTOF SCP system. In line with the previous results, we observed a signal increase when using AA, leading to increased proteome coverage at every peptide load assessed. Relative improvements were more apparent at lower concentrations, with a 20 picogram peptide digest demonstrating a striking 1.8-fold increase to over 2,000 protein groups identified in a 30 minute analysis. Furthermore, we find that this increase in signal can be leveraged to reduce ramp times, leading to 1.7x more scans across each peak and improvements in quantification as measured by %CVs. When evaluating single cancer cells, approximately 13% more peptide groups were identified on average when employing AA in the place of FA. All vendor raw and processed data are available through ProteomeXchange as PXD046002 and PXD051590.

**TOC Graphic:** 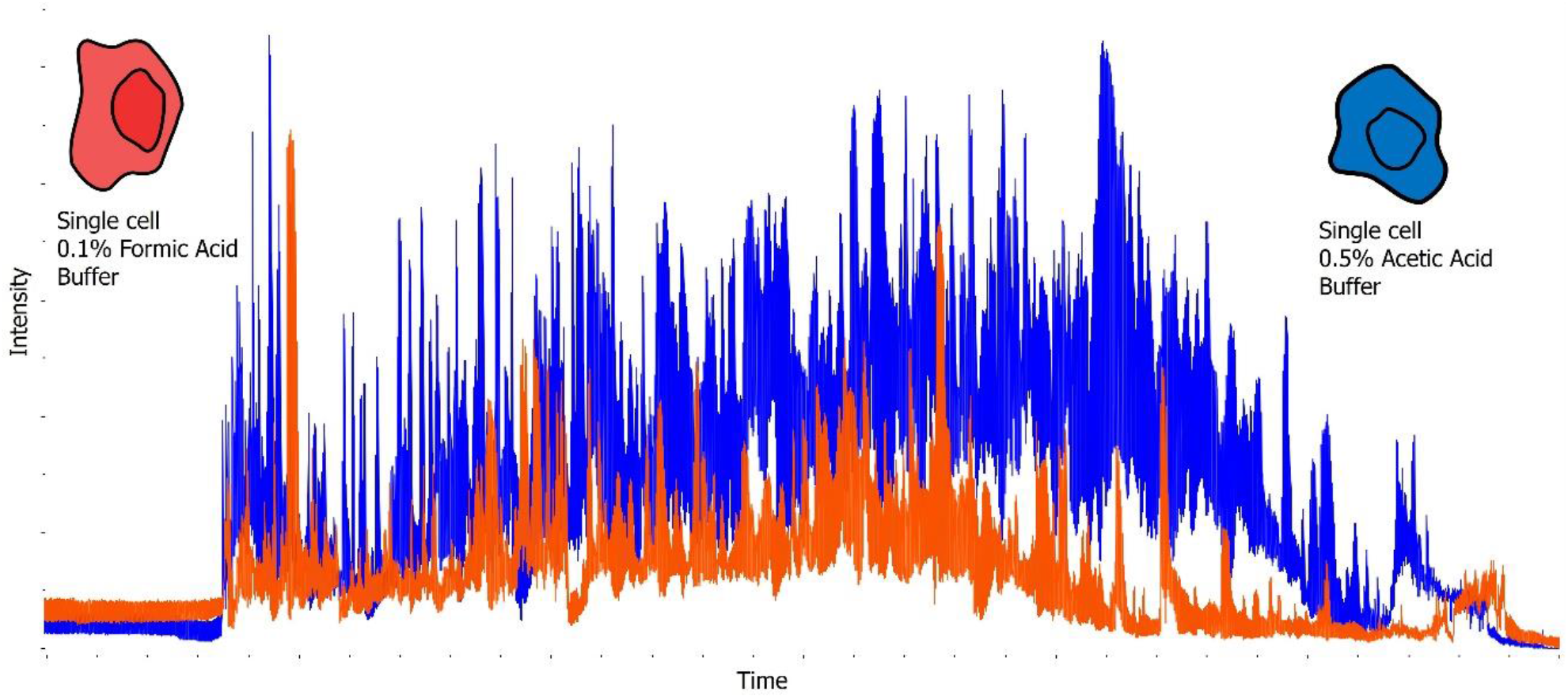

## Introduction

Single cell proteomics (SCP) by liquid chromatography-mass spectrometry (LCMS) is a growing field of research driven by rapid innovation in sample preparation^1,2^, chromatography^3^, mass spectrometry methods^4–6^, instrumentation^7^ and data analysis.^8–10^ Despite these advances, improvements in peptide detection efficiency are still essential, as it has become a critical parameter when working with a finite amount of material in an extremely complex form. Recently, Battellino *et al*. demonstrated that substituting 0.1% formic acid (FA) for 0.5% acetic acid (AA) in their LCMS proteomics workflow increased peptide signal 2.5-fold, with only slight decreases in chromatographic resolution.^11^ While the use of 0.5% AA may provide negligible benefits in shotgun proteomics workflows where sample is typically less sparse, this change in buffer could result in a remarkable improvement in SCP workflows.

## Methods

### Preparation of K562 dilution series

Promega human cancer cell line tryptic digest (V7461) was serial diluted using a solution containing 0.1% FA in LCMS grade water (Pierce 85170) and 0.1% n-Dodecyl-beta-Maltoside Detergent (DDM, Thermo Fisher, 89902). A total of 10 microliters of each dilution, 10 pg/μL, 100 pg/μL, and 1000 pg/μL, were loaded into 16 wells each of a 96 well plate. The plate was tightly sealed with plate sealing adhesive tape (Fisher, 60180-M143) and centrifuged prior to loading on the autosampler.

### Isolation and preparation of cancer cells

Cancer cell lines were obtained from ATCC and grown in the appropriate culture media described by the vendor. PANC 0203 cells were grown in RPMI 1640 (ATCC 30-2001) supplemented with 15% fetal bovine serum (FBS) (ATCC 30-2020) and 10 units of human insulin (Fisher). SW620 cells were grown in Leibovitz’s L-15 Medium (ATCC 30-2008) supplemented with 10% FBS (ATCC 30-2020). NCI-H-358 cells were grown in RPMI 1640 (ATCC 30-2001) supplemented with 10% FBS (ATCC 30-2020). All culture media was supplemented with 10 mg/mL Penn Strep antibiotic solution (ATCC 30-2300). All cell lines were passaged a minimum of 3 times prior to single cell isolation. Cells were harvested first by vacuum aspiration of the cell culture media. The adherent cells were briefly rinsed in 3 mL of 0.05% Trypsin plus EDTA solution (ATCC 30-2001). This solution was rapidly aspirated off and replaced with 3 mL of the same solution. The cells were examined by light field microscopy and incubated at 37°C with multiple examinations until the adherent cells had lifted off the plate surface, approximately 5 minutes. The active trypsin was then quenched by the addition of 7 mL of the original culture media. The 10 mL solution was transferred to sterile 15 mL Falcon tubes (Fisher) and centrifuged at 300 *x g* for 3 minutes to pellet the cells. The supernatant was gently aspirated off and the cells were resuspended in PBS solution without calcium or magnesium with 0.1% BSA (both, Fisher Scientific) at a concentration of 1 million cells per mL, as estimated by bright field microscopy. Cells for single cell aliquoting were gently dissociated from clumps by slowly pipetting a solution of approximately 1 million cells through a Falcon cell strainer (Fisher, 353420). The cells were placed on wet ice and immediately transported to the Johns Hopkins University Bloomberg Flow Cytometry and Immunology Core. Non-viable cells were labeled with a propidium iodide solution. The cell suspension was briefly vortexed prior to cell isolation and aliquoting.

Single cells were aliquoted using an analog MoFlo sorter into cold 96-well plates containing 2 μL of LCMS grade acetonitrile (one cell per well). Following aliquoting, each plate was immediately sealed and placed in an insulated box of dry ice with the wells pressed into the material to ensure rapid cooling. The plates containing frozen single cells were stored at -80°C (PANC 0203 and H-358) or processed immediately (SW620). For processing, acetonitrile was driven off by heating the cells for 90 seconds at 95°C using a 96-well hotplate. Dried cell lysates were digested using 2 μL of a digestion solution containing 5 ng/μL LCMS grade trypsin (Pierce) in 0.1% DDM (Thermo Fisher, 89902) and 50mM TEAB (Thermo Scientific). The plates were sealed with adhesive plate tape (Fisher, 60180-M143), and digestion occurred at room temperature overnight. Following digestion, the plates were briefly centrifuged to condense evaporation and wells were completely dried under vacuum centrifugation (Eppendorf Vacufuge Plus). Resulting peptides were resuspended in 3.5 μL of 0.1% FA, vortexed and centrifuged prior to loading on the autosampler.

### LCMS instrument parameters

An EasyNLC 1200 system (Proxeon) coupled to a TIMSTOF SCP (Bruker Daltronic) was used for all analyses. Peptides were separated using a constant flow of 300 nL/min on an IonOpticks 15 cm x 75 μm C-18 column with 1.5 μm particle size (Ion Opticks “Ultimate”) with integrated CaptiveSpray emitter held at 1500V. For standard injections, an appropriate volume (1-4 μL) of each well was loaded with a partial loop injection of 7 - 10 μL at 900 bar. Pickup and partial loop injection volume values (2 x pickup volume + 2 μL if > 7 μL) were used according to the recommendations of the “How to set up the EasyNLC method” document by the University of Washington Proteome Research Center (https://proteomicsresource.washington.edu/). For single cell injections, 4 μL was picked up with a partial loop injection volume of 10 μL. Buffer A consisted of either 0.1% FA (Thermo Scientific) or 0.5% AA (Fisher Scientific) in water (Fisher Chemicals). Buffer B consisted of 80% acetonitrile in water with the appropriate ion pairing modifier for each experiment. All reagents were LCMS grade. The 30 minute gradient used in all experiments began at 8% buffer B and ramped to 35% B by 22 minutes. The gradient then increased to 100% B by 26 minutes where it held for 2 minutes before returning to baseline conditions. The column was equilibrated in 12 microliters of baseline buffer conditions prior to each injection.

The TIMSTOF SCP system was operated in diaPASEF mode using a method with 50 Da isolation windows provided by Dr. Michael Krawitzky of Bruker Daltronic during instrument training and will be referenced as “default parameters” in the study. In this method, the MS1 scanned from 100-1700 m/z with a 1/k0 window of 0.6-1.4. Ions that entered the mass analyzer were restricted by a user defined polygon set on the m/z vs 1/k0 heatmap. The polygon can impart dramatic alterations on ion signal, thus must be carefully optimized for each series of experiments. In this experiment, the polygon began at 300 m/z and was set to not acquire ions in the MS1 cloud below 800 m/z. The method utilized 6 cycles with 3 isolation windows per cycle, resulting in a total time of 1.2 seconds when a 166 ms ramp time was used. The “high sensitivity” mode was enabled for all samples. For the 100 ms experiments described, the ramp time was decreased from 166 ms to 100 ms to allow a shorter cycle time of 0.72 seconds. To prepare the instrument for samples utilizing a different ion pairing modifier, buffer bottles were changed, and 5 full purge cycles were performed. The precolumn and analytical columns were equilibrated using 10 μL and 12 μL of mobile phase A, respectively, at 900 bar to flush the previous ion pairing modifier from the columns.

### Data Analysis

Prior to analysis, all raw files were converted to the HTRMS file format using the “HTRMS Converter” software (Biognosys) using default parameters. These files were then analyzed in SpectroNaut 18 (Biognosys) via the directDIA+ workflow with default parameters for data calibration and analysis. The UniProt SwissProt reviewed library for human (downloaded on March 3, 2023) and appended with the cRAP contaminant database (www.gpm.org) was used for library generation. For standard digests carbamidomethylation of cysteines were considered as static and methionine oxidation was considered a dynamic modification. Single cells were analyzed exclusively with methionine oxidation. Output data was visualized in GraphPad Prism 10.0.1. All Bruker .d raw files and SpectroNaut processed results have been made publicly available through the ProteomeXchange partner repositories^12^ as PXD046002 and PXD051590.

## Results

### AA improves proteomic coverage on sub-nanogram injections of peptides

To determine whether the use of FA or AA in mobile phase impacts proteomic coverage, a dilution series of a standard trypsin digest was injected into the instrument with 20, 100, 200, 400 and 1,000 pg total peptide on column using both buffer conditions. Our data show an increase in detected precursors, peptides, proteins, and protein groups in the 20-200 pg dilutions. Interestingly, as peptide concentration on column increased from 20-200 pg, the number of additional identifications gained by AA modifier compared to FA modifier decreased. Remarkably, at 20 picogram of digest standard the number of unique peptides detected nearly doubled, leading to the striking observation of 2,000 proteins groups at this concentration (**Figure 1**). Of note, at 200 picograms or above of standard trypsin digest on column, the number of precursors and peptide groups were higher in the AA buffer but did not translate to a significant increase in protein groups detected (**Supplemental File 1**).

**Figure 1.**
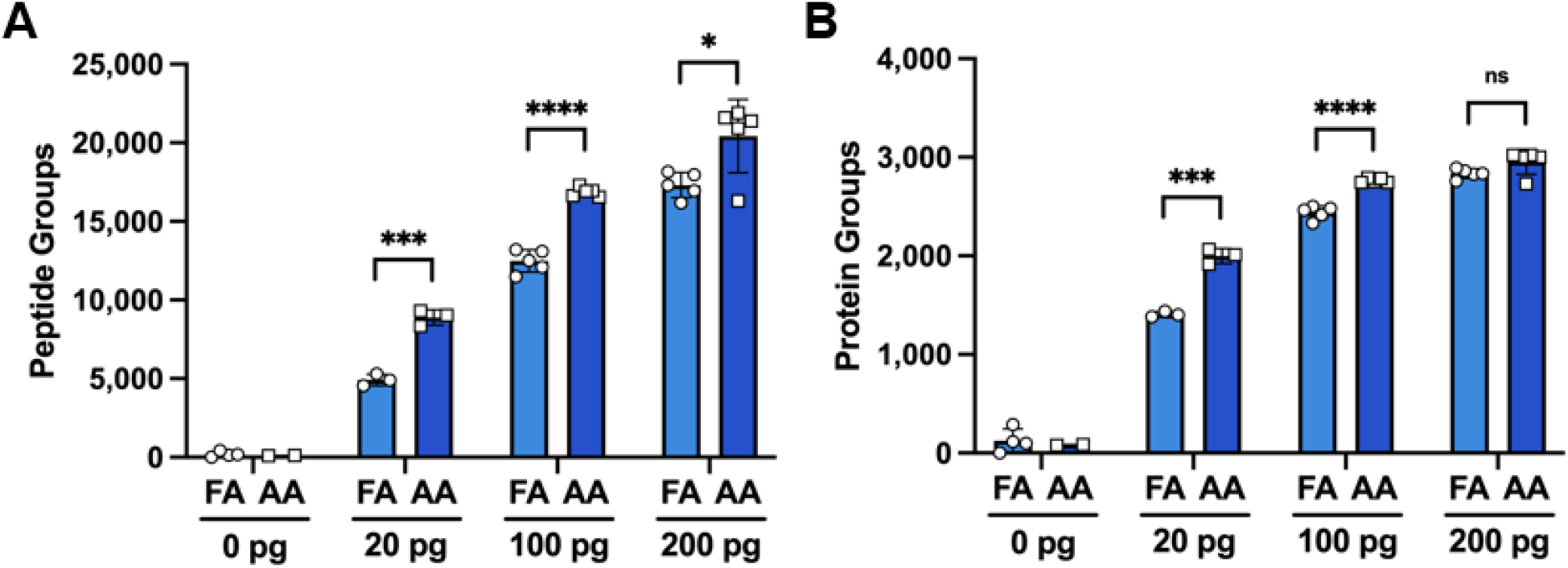
Proteomic coverage of sub-nanogram injections of a standard trypsin digest. (**A**) Peptide and **(B)** protein groups identified at 20, 100, and 200 pg of peptide digest with 0.1% formic acid (FA) or 0.5% acetic acid (AA) in the mobile phases. Error bars represent mean and standard deviation. Statistical analyses were performed using a two-tailed t-test. P-value < 0.05*, < 0.001 ***, < 0.0001****; ns: no significance.

### Increased relative signal from AA buffer can be used to obtain more scans across each peak

In addition to an increase in the number of peptide and protein groups identified, the overall signal intensity was also higher when using AA as the ion pairing modifier. We hypothesized that the increased relative signal provided by AA could be exploited to reduce relative ion accumulation time, resulting in more scans across each peak. The default ramp time on the TIMSTOF SCP system and vendor provided methods is 166 ms, resulting in a cycle time of 1.2 seconds. If the ramp time is decreased to 100 ms, the instrument software estimates a cycle time of 0.7 seconds, allowing 1.7 times more scans across each peak. In general, more measurements allow improved relative quantification, which is typically assessed in shotgun proteomics by calculating the coefficient of variation (%CV).^13^ We next repeated analysis of the digest dilution series using the 100 ms ramp time with the AA buffer system. The resulting number of peptides with %CVs below 20% and 10% were then extracted from the SpectroNaut post-analysis report (**Figure 2**). We observed that in the 100 pg and 200 pg standard trypsin digests the number of peptides with low CVs increased with the substitution of AA and, further, with the decrease in ramp time. However, the 20 pg dilution in AA buffer with a 100 ms ramp time exhibited a decrease in the quantitative accuracy when compared to the 166 ms ramp time.

**Figure 2.**
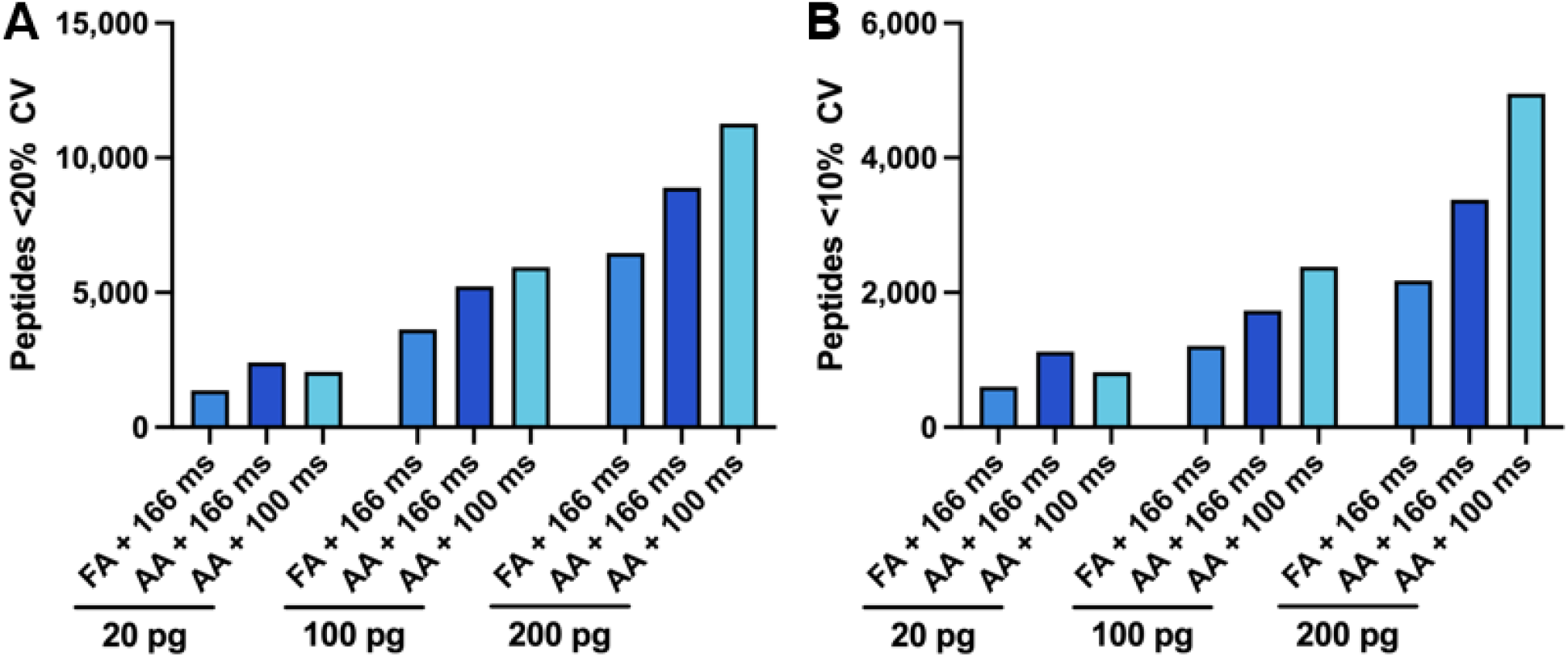
The number of peptides quantified with low %CVs. Peptides quantified at less than **(A)** 20% CV and **(B)** 10% CV when injecting 20, 100, and 200 pg of standard trypsin digests with 0.1% formic acid (FA) or 0.5% acetic acid (AA) buffers with ramp times 166 ms or 100 ms.

### AA increases protein sequence coverage in isolated single human cells

Encouraged by these results, we repeated this analysis using isolated single human cancer cells from three separate cell lines. Remarkably, using this extremely simple single cell preparation method, over 800 protein groups were identified on average across the 98 single cells which returned quantifiable data (**Supplemental File 1**). While protein group numbers this high have been reported by others, careful monitoring of sample loss with picoliter robotics^1,5^ or other advanced systems^14^ has been required. In evaluating the impact of ion pairing modifier on single cell proteomics, we observed a general upward trend in both peptide and protein groups identified when employing AA (**Figure 3, Supplemental Figure 1**). Of particular interest, single H-358 cells were analyzed with both the default 166 ms and reduced 100 ms ramp time methods. When utilizing a ramp time of 166 ms, an improvement of approximately 14% was observed in the number of peptides and precursors identified for single cells run using AA buffers **(Figure 3A**). It should be noted that a two-tailed t-test still found this increase to be significant even when the two lowest cells analyzed with FA buffer were removed from consideration (data not shown). However, the increase in the number of peptides did not lead to a significant increase in the number of protein groups identified (**Figure 3B**). When utilizing the 100 ms ramp time, both a higher number of peptides and proteins were observed when AA was used as a modifier, but in neither case was this improvement found to be statistically significant. Similar to the 20 pg peptide standard the number of peptide and protein groups was lower when the 100 ms ramp time was used.

**Figure 3.**
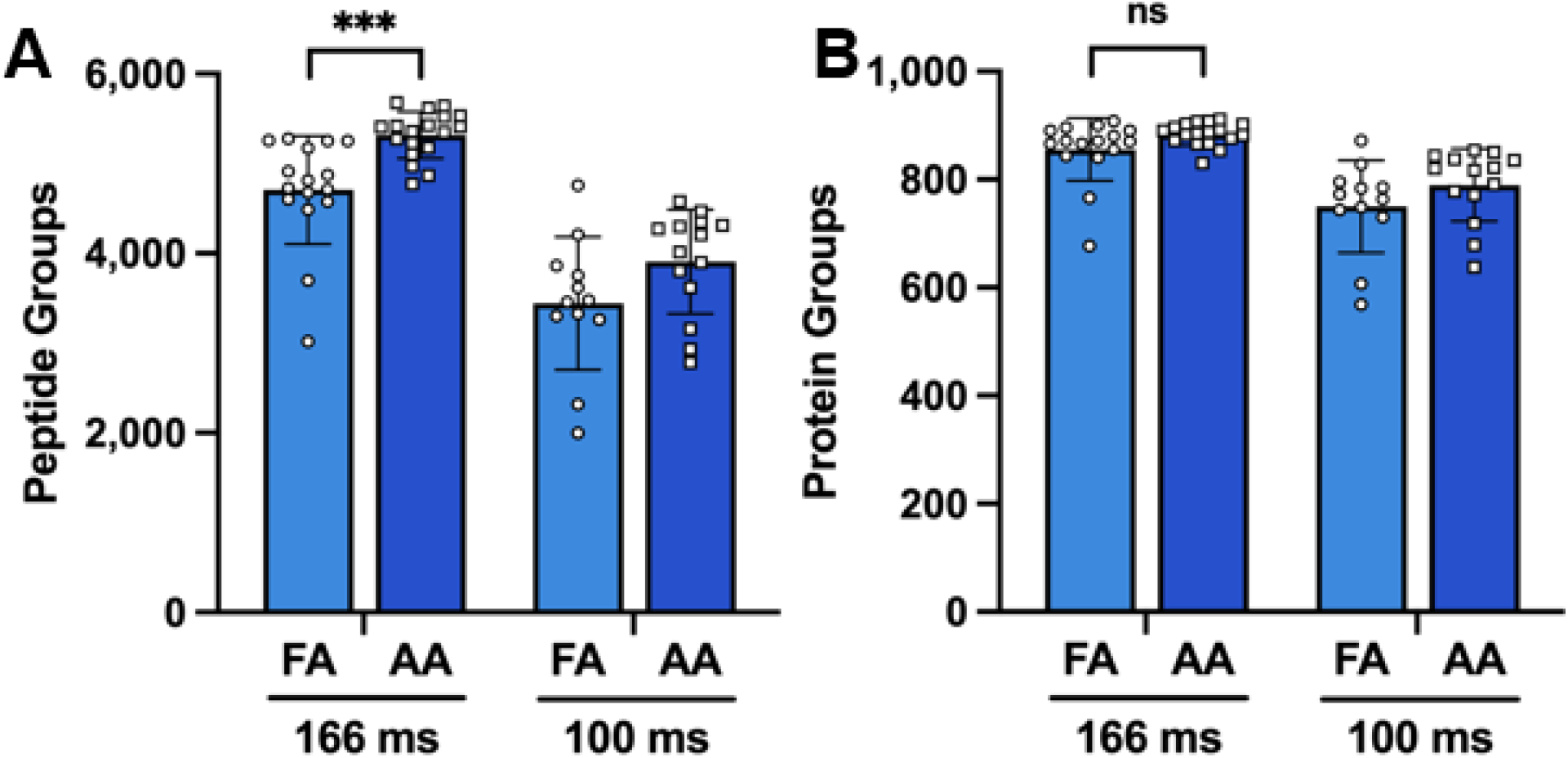
Peptide and protein groups identified in single cells. (**A**) Peptide and **(B)** protein groups identified in single H-358 cancer cells with 0.1% formic acid (FA) or 0.5% acetic acid (AA) in the mobile phases when employing ramp times of 166 ms and 100 ms. Error bars represent mean and standard deviation. Statistical analyses were performed using a two-tailed t-test. P-value < 0.001 ***, ns: no significance.

## Conclusions

Adulterating running buffers to increase peptide signal and identification rates is not a new concept. The addition of 5% DMSO has long been noted as an additive that can lead to impressive increases in peptide and protein coverage relative to 0.1% FA alone.^15^ As noted in Battellino *et. al*., however, less than 2% of all proteomics studies in the literature have employed DMSO.^11^ We suspect this is due to reports of contamination issues after long term use which can require extensive cleaning of internal instrument components. AA appears to provide comparable increases in peptide signal and has been employed in LCMS systems used for the analysis of extractables and leachables and pesticides for decades.^16,17^ It seems unlikely, for this reason, to be detrimental to long term instrument performance. With similar increases in total ion signal, AA is an attractive alternative and appears applicable across a wide range of gradients and instrument conditions. In Battellino *et al*., gradients between 60 and 120 minutes in length were effective for proteomics on both QTOF and Orbitrap analyzers. Here we have found similar results for 30 minute gradients on a specialized TIMSTOF system designed for low input samples.

While it is curious that we see less of an improvement in coverage when analyzing isolated single human cells, we observe a general trend of improvements in three separate preparations of single cells. The evaluation of this disconnect is outside of the scope of this study but could simply be due to sampling bias as only 98 cells were successfully analyzed across three separate cell lines and two different instrument methods. We can, however, make some inferences here on the amount of sample loss in this simple one pot sample preparation method. Work in our group previously estimated each H-358 cell grown in these culture conditions to have an average protein content of 200 pg.^18^ At a peptide group level, we observe approximately the same number of peptides as the 20 pg standard digest, though lower overall protein numbers. Similarly, we find that decreasing the TIMS ramp time to 100 ms from the default 166 ms method results in a loss in proteome coverage in both single cells and 20 pg of standard digest. Taken together, these suggest that this single cell preparation method results in a 90% loss of protein content. This may not be surprising given the relatively large surface area available for peptide adhesion within a standard 96-well plate, compared to loss inferences from cells prepared directly within 384-well plates in recent studies.^19–21^ Our interpretation is that the use of highly diluted peptide standards as a proxy for isolated single cells, while seemingly practical, may produce results that should be interpreted with a high degree of caution. However, the final conclusion of this technical note is that the emerging field of single cell proteomics should continue to question traditional proteomics methods at every stage, as remarkable increases in coverage appear to be achievable through relatively minor optimizations.

## Acknowledgements

We would like to thank Ahmed Warshanna and Hannah Wilkins for helpful conversations and technical assistance with various aspects of this study.

---

**Supplemental Figure 1.**
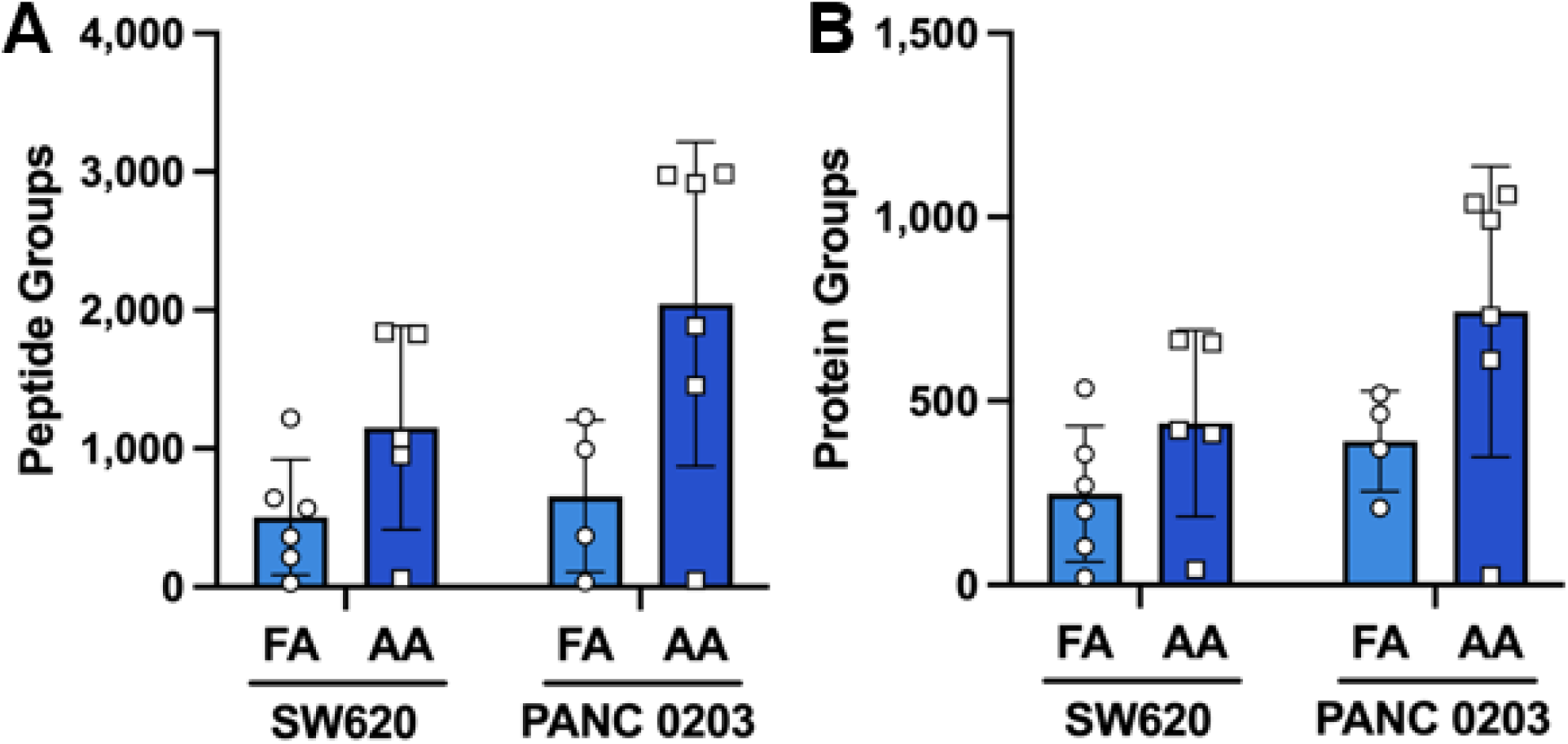
Peptides and proteins obtained from single cells. **(A)** Peptide and **(B)** protein groups identified in SW620 and PANC 0203 cells analyzed using a single step digestion method and run using 0.1% formic acid (FA) or 0.5% acetic acid (AA) buffer additives. Error bars represent mean and standard deviation. Statistical analyses were performed using a two-tailed t-test and FA vs AA comparisons resulted in no significant differences.

